# Discovery of a new *Myristica* swamp in the Northern Western Ghats of India

**DOI:** 10.1101/2024.02.12.579903

**Authors:** Pravin Desai, Vishal Sadekar, Shital Desai

## Abstract

*Myristica* swamps are one of the world’s unique freshwater ecosystems. In recent years there has been an increase in the number of reports about their distribution along the western Ghats. In the current paper, we report a new distribution record for the occurrence of *Myristica* Swamp from the northern part of the western Ghats of Maharastra. The *Myristica* swamp is located within the Bhalandeshwar Sacred Grove of Kumbral Bagwadi, Dodamarg Maharastra, India. This forms a second report from the state of Maharastra. Frequent field visits were made to study and document the floral assemblage and biodiversity in the swamp from, November 2023 to January 2024. The swamp is dominated by *Myristica magnifica* Bedd. The swamp is a part of a sacred grove and it is protected by Villagers due to religious beliefs. The discovery of this swamp points towards the possibility of the occurrence of more swamps in this region. Hence there is a need for a systematic survey for documenting swamps in the different areas.

---

*Myristica* swamps represent highly important yet one of the threatened freshwater ecosystems of the world. As the name suggests, the plants of the family Myristicaceae dominate the swamps. The high watershed value of these swamps supports many rare, endemic flora and fauna (Chandran et al. 2010). They are also denoted as living fossils due to the primitive nature of *Myristica* plants (Roby et al. 2013). With an evolutionary origin of about 140 million years, the swamps are valuable for evolutionary studies (Chandran et al. 1999; Dharmapalan and Asokhan 2013).

Krishnamoorthy (1960) for the first time, shed light on the occurrence of this unique ecosystem from the south Western Ghats of Kerala, India. Since then, their occurrence has also been studied and reported in the states of Karnataka, Goa, and Maharashtra (Gadgil & Chandran 1989; Santhakumaran et al. 1995; Jose et al. 2014; Sreedharan and Indulkar 2018). The *Myristica* swamp reported from Bambarde-Hewale of Maharashtra forms the first record of the presence of such an ecosystem in the state and the northernmost record in the Western Ghats (Sreedharan and Indulkar 2018).

The literature survey indicates that the swamps show restricted distribution along the Western Ghats of India in small fragmented areas. In the present communication, we are reporting a new site occurrence of *Myristica* swamps from Maharashtra state, India. Pravin Desai runs a homestay and conducts wildlife trails in the different parts of Dodamarg Tilari Bioregion in Dodamarg Taluka of Sindhudurg District of Maharashtra. During his regular birding trail to Bhalandeshwar sacred grove, while searching for Brown Wood-owls (*Strix leptogrammica*) he observed a huge tree of *Myristica magnifica* Bedd. which led to him discovering the *Myristica* swamp. This was subsequently followed by our visits to the swamp for further studies.

We estimated the area of the sacred grove and the area of the swamp within the sacred grove using Garmin GPS 72s by walking around the grove and swamp. We measured all trees in the swamp with≥ 30 cm girth at breast height by using a measuring tape, and height by using a Leica geosystem D1 distometer. For *Myristica magnifica*, an obligate swamp specialist, we recorded the height and girth of all individuals. For individuals who were less than 1.3 cm in height, their girth was recorded (ocular estimation). Tree seedlings (GBH < 10 cm) were enumerated from the sample plots to determine the regeneration status of tree species in the *Myristica* swamps. Additionally, we have created a checklist for other woody plant species to understand the species diversity and identify the associated species in the area.

The *Myristica* swamp is located in the Bhalandeshwar sacred grove at Kumbral Bagwadi (Fig. 1). The swamp is dominated by *Myristica magnifica*, which has prominent stilt roots Image 1. The area of the grove and swamp is 8200 m^2^ and 770 m^2^ respectively. A total of 39 plant species were documented (Table 1). We recorded 70 individuals of *Myristica magnifica*, out of which 19 are with a girth of ≥ 30 cm and 51 individuals with a girth of < 30 cm. Given that 51 out of 70 individuals were < 30 cm indicates regeneration of the *Myristica magnifica*. The size class plot for girth and height is given in Fig. 2 and 3 respectively. The rank abundance curve shows that the swamp species exhibit low species evenness, with *Myristica magnifica* being the dominant species in the swamp (Fig. 4). The undergrowth in the swamp is dominated by the fern *Bolbitis presiliana* Ching and *Pandanus furcatus* Roxb (Image 1).

**Table 1.** Checklist of plants documented in the Myristica swamp of Bhalandeshwar.

| Sr.no | Species Name | Family | Habit | IUCN status |
| --- | --- | --- | --- | --- |
| 1 | *Actephila excelsa* (Dalzell)<br>Müll.Arg. | Phyllanthaceae | Small tree | Least Concern |
| 2 | *Allophylus cobbe* (L.) Forsyth f. | Sapindaceae | Liana | NA |
| 3 | *Anamirta cocculus* (L.) Wight & Arn. | Menispermaceae | Liana | NA |
| 4 | *Antidesma nigricans* Tul. | Phyllanthaceae | Shrub | NA |
| 5 | *Aporosa cardiosperma* (Gaertn.) Merr. | Phyllanthaceae | Tree | Vulnerable |
| 6 | *Artocarpus hirsutus* Lam. | Moraceae | Tree | Least Concern |
| 7 | *Bolbitis presiliana* Ching. | Dryopteridaceae | Tree | Least Concern |
| 8 | *Bridelia retusa* (L.) A.Juss. | Phyllanthaceae | Tree | Least Concern |
| 9 | *Bridelia scandens* | Phyllanthaceae | Scandent shrubs | Least Concern |
| 10 | *Calophyllum apetalum* Willd. | Calophyllaceae | Tree | Vulnerable/ Endemic |
| 11 | *Capparis spp* | Capparaceae | Climber | NA |
| 12 | *Carallia brachiata* (Lour.) Merr. | Rhizophoraceae | Tree | Least Concern |
| 13 | *Caryota urens* L. | Arecaceae | Tree | Least Concern |
| 14 | *Cleodendron sp.* | Lamiaceae | Shrub | NA |
| 15 | *Combretum sp.* | Combretaceae | Herb | NA |
| 16 | *Diospyros thwaitesii* (Hiern) Bedd. | Ebenaceae | Tree | Vulnerable |
| 17 | *Dracaena elliptica* Thunb. & Dalm. | Asparagaceae | Herb | Least Concern |
| 18 | *Ficus hispida* L.f. | Moraceae | Tree | Least Concern |
| 19 | *Ficus nervosa* Roth | Moraceae | Tree | Least Concern |
| 20 | *Garcinia indica* (Thouars) Choisy | Clusiaceae | Tree | Vulnerable |
| 21 | *Gymnosporia rothiana* (Walp.) M.A.Lawson | Celastraceae | Shrub | NA |
| 22 | *Holigarna arnottiana* Hook.f. | Anacardiaceae | Tree | Endemic |
| 23 | *Ipomoea campanulata* L. | Convolvulaceae | Climber | NA |
| 24 | *Ixora nigricans* R.Br. ex Wight & Arn. | Rubiaceae | Small tree | NA |
| 25 | *Ixora brachiata* Rox. | Rubiaceae | Tree | NA |
| 26 | *Lagenandra toxicaria* Dalzell | Araceae | Herb | Least Concern |
| 27 | *Leea indica* (Burm.f.) Merr. | Vitaceae | Small tree | Least Concern |
| 28 | *Lophopetalum wightianum* Arn. | Celastraceae | Tree | Least Concern |
| 29 | *Macaranga peltata* (Roxb.) Müll.Arg. | Euphorbiaceae | Tree | NA |
| 30 | *Machilus glaucescens* (Nees) Wight | Lauraceae | Tree | NA |
| 31 | *Myristica magnifica* Bedd. | Myristicaceae | Tree | Endangered/<br>Endemic |
| 32 | *Nothopegia castaneifolia* (Roth) Ding Hou | Anacardiaceae | Tree | Critically Endangered<br>/ Endemic |
| 33 | *Pandanus furcatus* Roxb | Pandanaceae | Shrub | NA |
| 34 | *Pothos scandens* L | Araceae | Climber | NA |
| 35 | *Premna coriacea* C.B.Clarke | Lamiaceae | Climber | NA |
| 36 | *Pterospermum acerifolium* (L.) Willd. | Malvaceae | Tree | Least Concern |
| 37 | *Sterculia guttata* Roxb | Malvaceae | Tree | NA |
| 38 | *Tabernaemontana alternifolia* L. | Apocynaceae | Tree | NA |
| 39 | *Tetrastigma* sp. | Vitaceae | Climber | NA |

**Figure 1.**
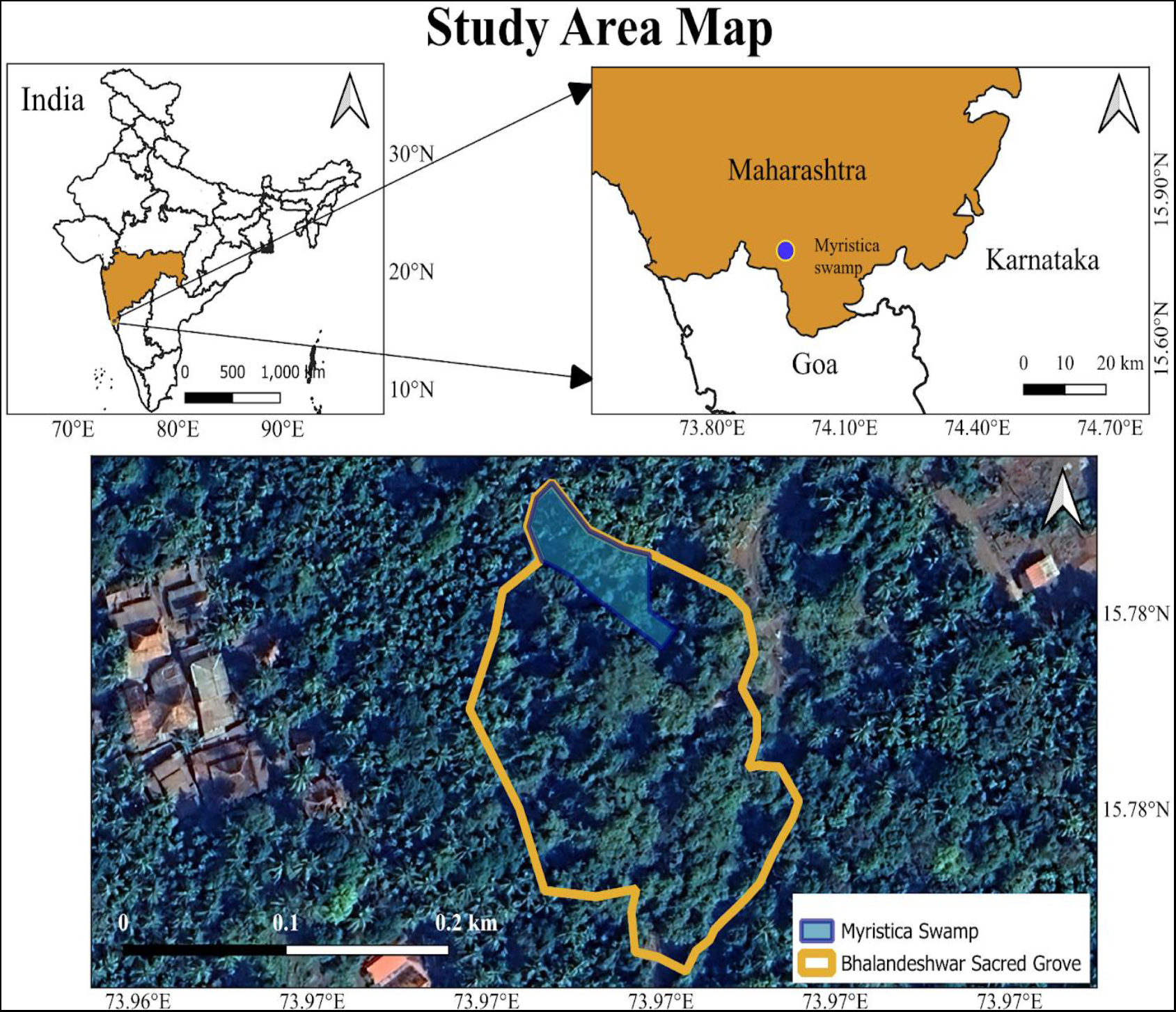
Map showing the location of Bhalandeshwar Sacred Grove and Myristica Swamp as on December 16, 2023.

**Figure 2.**
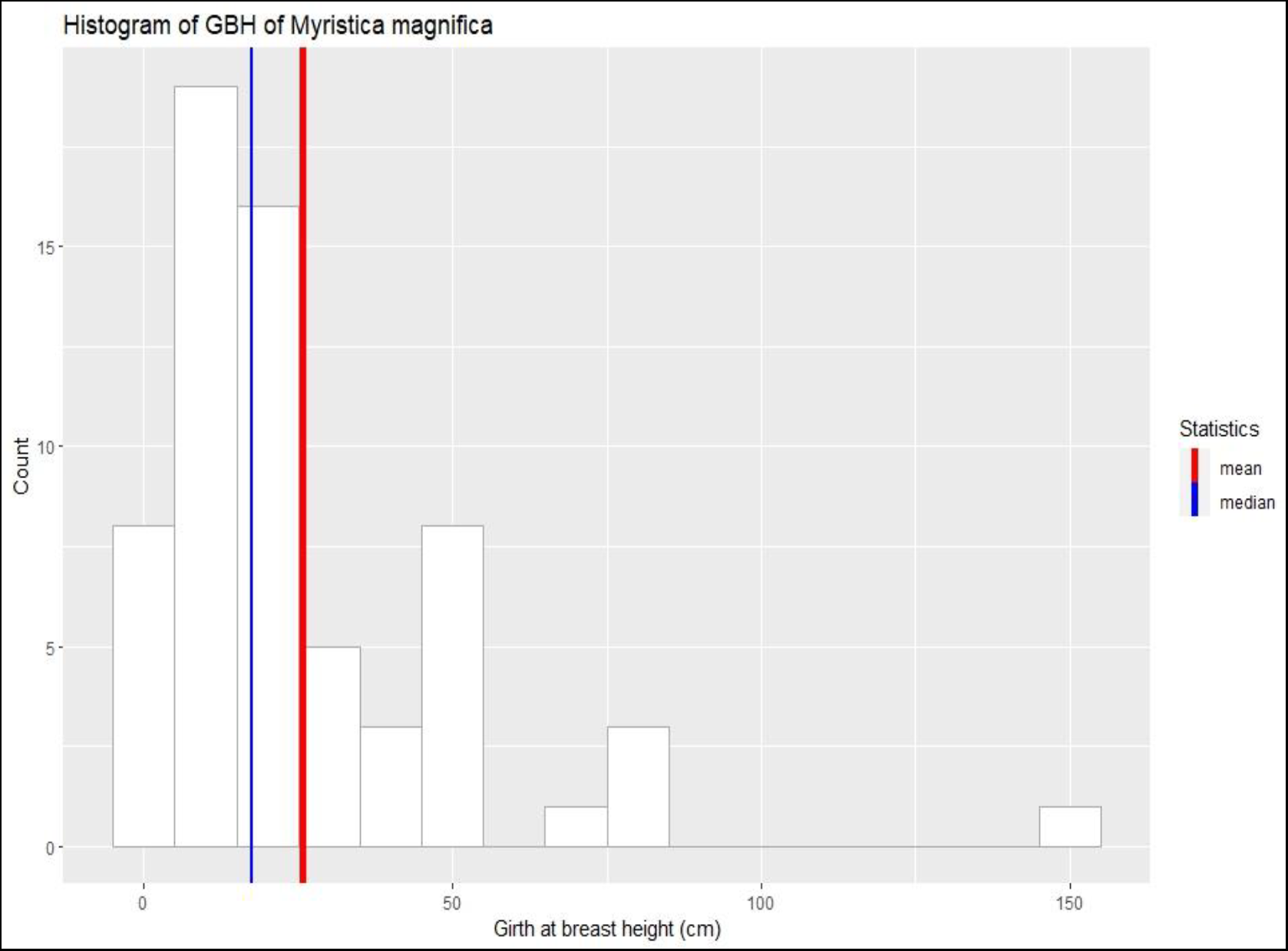
Histogram of girth at breast height of *Myristica magnifica* individuals that were > 1.3 cm in height.

**Figure 3.**
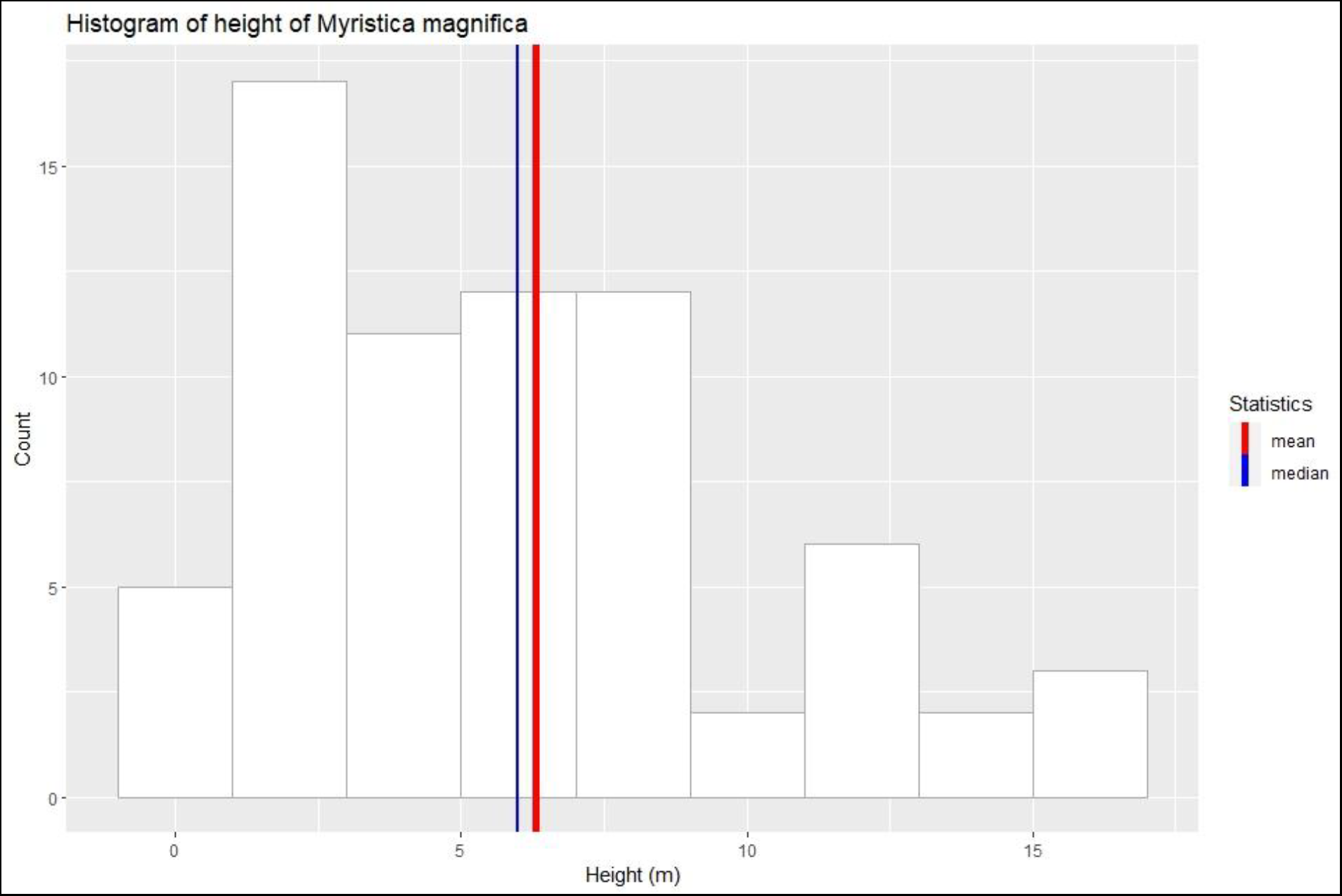
Histogram showing *Myristica magnifica* tree height distributions in the swamp. There is a greater representation of saplings indicating the regeneration of trees in the swamp.

**Figure 4.**
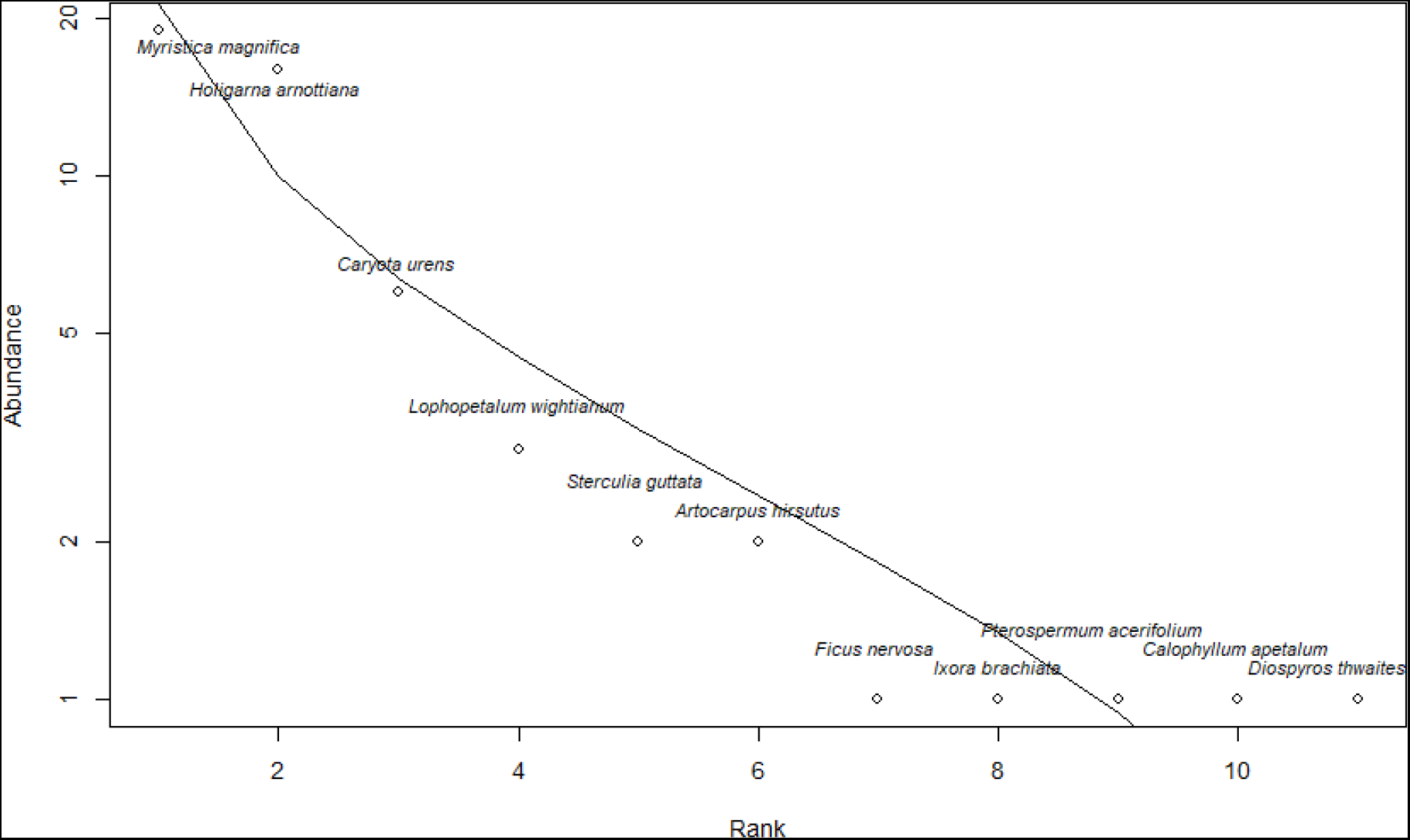
Rank abundance curve for all individual woody plant tree species in the *Myristica* swamp. Only those individuals with ≥30 cm girth at breast height were included in this plot.

The local communities worship the deity Bhalandeshwar, who is believed to be an avatar of Lord Shiva and they have been performing religious rituals since the 16^th^ century. According to the elders, the land was awarded as jagir during the reign of the Adilshahi Dynasty. The spring that emerges at the temple, serves as a source of drinking water for local people. The swamps offer various ecological services, like groundwater recharge, carbon sequestration, natural barriers against floods, habitat, and food for many aquatic and aerial fauna. The fruits of *Myristica* are important food plants for threatened hornbills (Gopal et al. 2021). The occurrence and discovery of this second swamp in the northern Western Ghats of Maharashtra strongly point toward the possibility of more swamps in this region. Hence, a systematic survey is needed to document swamps in the different areas. Religious beliefs and sustainable use of the water from swamps have been the important reasons for protecting this swamp for several decades.

## Acknowledgments

The authors thank Zilba Desai for providing cultural and traditional information about the sacred grove and swamp and Raman Kulkarni, Honorable Wildlife Warden of Kolhapur, for the photographs. We also thank Rohit Naniwadekar, Himanshu Lad, Prashant Jadhav, and Faruk Mehtar for their guidance and Navendu Page for help with identification. All the villagers of Kumbral are also acknowledged for conserving the swamp.

---

**Image 1.**
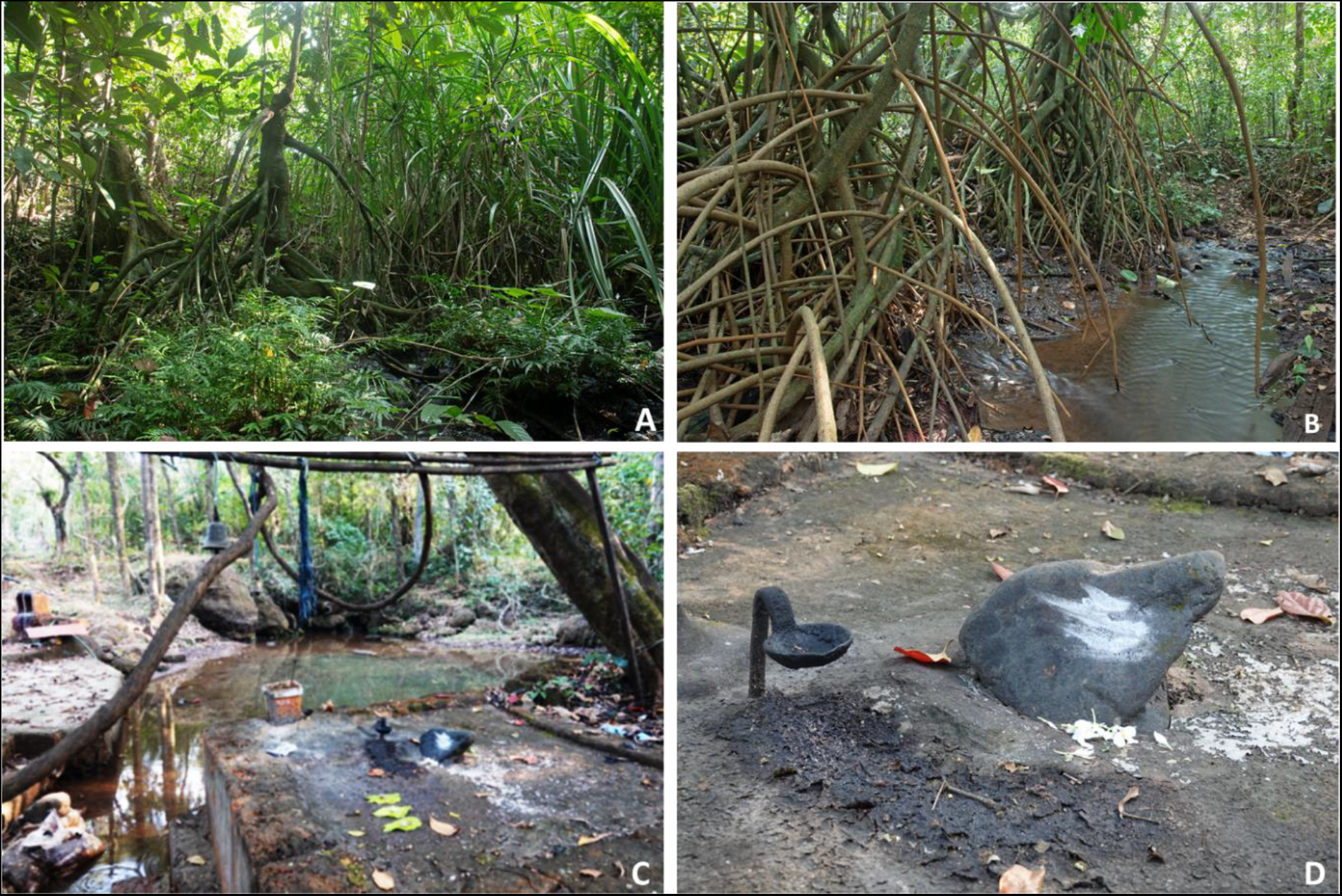
A) *Myristica magnifica* with undergrowth of fern, *Bolbitis presiliana*, and edge vegetation B) Water inundation in the swamp and Stilt roots of *Myristica magnific*a C) Source of water for the swamp D) God idol being worshiped in the sacred grove.

## Author Contributions

Pravin Desai discovered the swamp and assisted with fieldwork; Vishal Sadekar conducted the field survey and wrote the manuscript; Shital Desai assisted with fieldwork and wrote the manuscript. All authors contributed critically to the drafts and gave final approval for publication.

## References

Chandran, M. D. S., Mesta, D. K., & Naik, M. B. (1999). Inventorying and conservation of the Myristica swamps of Uttara Kannada. Report of Research and Training Institute, Bangalore.

Chandran, M. S., Rao, G. R., Gururaja, K. V., & T.V Ramachandra. (2010). Ecology of the swampy relic forests of Kathalekan from Central Western Ghats, India. Bioremediation, Biodiversity and Bioavailability 4(1): 54–68

Dharmapalan, B. & Asokhan A (2013). Myristica swamps–evolutionary relics. Science Reporter, June 2013, pp 45–48

Gadgil, M. & Chandran, M.D.S. (1989). Environmental Impact of Forest Based Industries on the Evergreen Forests of Uttara Kannada District, A Case Study (Final Report). Department of Ecology and Environment, Bangalore. 20 pp

Gopal, A., Mudappa, D., Raman, T.R.S., & Naniwadekar, R. (2021). Seed fates of four rainforest tree species in the fragmented forests of Anamalais in the southern Western Ghats, India. Acta Oecologica, 110, 103698.

Jose, J., Roby, T. J., Ramachandran, K. K., & Nair, P. V. (2014). Species abundance distributions of selected communities in the Myristica swamp forests of southern Kerala. Current Science, 447–453.

Krishnamoorthy, K. (1960). Myristica swamps in the evergreen forests of Travancore. Indian Forester 86(5): 314–315.

Roby, T. J., Jose, J., & Nair, P. V. (2013). Syzygium travancoricum (Gamble)–a critically endangered and endemic tree from Kerala, India–threats, conservation, and prediction of potential areas; with special emphasis on Myristica swamps as a prime habitat. Int. J. Sci. Environ. Technol, 2(6), 1335–1352.

Santhakumaran, L. N., Singh, A., & Thomas, V. T. (1995). Description of a sacred grove in Goa (India), with notes on the unusual aerial roots produced by its vegetation. Wood. Oct-Dec, 24–28.

Sreedharan, G., & Indulkar, M. (2018). New distributional record of the northernmost Myristica swamp from the Western Ghats of Maharashtra. Current Science, 115(8), 1434–1436.

